# Modulation of Resting Connectivity Between the Mesial Frontal Cortex and Basal Ganglia

**DOI:** 10.1101/432609

**Authors:** Traian Popa, Laurel S. Morris, Rachel Hunt, Zhi-De Deng, Silvina Horovitz, Karin Mente, Hitoshi Shitara, Kwangyeol Baek, Mark Hallett, Valerie Voon

## Abstract

The mesial prefrontal cortex, cingulate cortex and the ventral striatum are key nodes of the human mesial fronto-striatal circuit involved in decision-making and executive function and pathological disorders. Here we ask whether deep wide-field repetitive transcranial magnetic stimulation (rTMS) targeting the mesial prefrontal cortex (MPFC) influences resting state functional connectivity. In Study 1, we examined functional connectivity using resting state multi-echo and independent components analysis in 154 healthy subjects to characterize default connectivity in the MPFC and mid-cingulate cortex (MCC). In Study 2, we used inhibitory, 1 Hz deep rTMS with the H7-coil targeting MPFC and dorsal anterior cingulate (dACC) in a separate group of 20 healthy volunteers and examined pre-and post-TMS functional connectivity using seed-based and independent components analysis. In Study 1, we show that MPFC and MCC have distinct patterns of functional connectivity with MPFC–ventral striatum showing negative, whereas MCC–ventral striatum showing positive functional connectivity. Low-frequency rTMS decreased functional connectivity of MPFC and dACC with the ventral striatum. We further showed enhanced connectivity between MCC and ventral striatum. These findings emphasize how deep inhibitory rTMS using the H7-coil can influence underlying network functional connectivity by decreasing connectivity of the targeted MPFC regions, thus potentially enhancing response inhibition and decreasing drug cue reactivity processes relevant to addictions. The unexpected finding of enhanced default connectivity between MCC and ventral striatum may be related to the decreased influence and connectivity between the MPFC and MCC. These findings are highly relevant to the treatment of disorders relying on the mesioprefrontal–cingulo–striatal circuit.

## I. Introduction

Neuromodulation with magnetic stimulation is emerging as a valuable treatment alternative for a wide range of psychiatric and neurologic disorders [1]. Repetitive transcranial magnetic stimulation (rTMS) is a technique that can be used to apply multiple brief magnetic pulses to neuronal structures, thus transiently modulating neural excitability in a manner that is dependent mainly on the intensity and frequency of stimulation [2]. It is a non-invasive, non-pharmacological, and safe treatment, in which abnormal communication within neuronal networks can be entrained and modified. Depending on the target, the depth at which stimulation occurs appears to be a crucial factor underlying potential therapeutic efficacy in certain disorders, such as major depressive disorder [3–5]. In this study, we investigate the modulation of resting neural activity in mesial prefrontal-striatal circuits in healthy subjects by inhibitory deep wide-field stimulation with an Hesed (H-)7 coil [6, 7].

Fronto-striatal circuits are critical for the processing of reward, anticipation of outcomes, and behavioral control [8–11]. Latent neural network organization and behavioral mechanisms in humans can be explored with resting state functional magnetic resonance imaging (fMRI) connectivity (rsFC), a method that measures the synchronization between intrinsic low-frequency fluctuations of brain regions in the absence of any specific task [12–14]. Since the connections identified at rest closely mirror anatomical connections [15] and predict brain activations associated with behavioral performance [16], rsFC is an important tool for characterizing *in vivo* circuit-level dynamics, which may support particular behavioral responses [17, 18].

Studies of substance use disorders have revealed the critical role of fronto-striatal circuits, highlighting large scale disruptions in functional connectivity between the mesolimbic reward system and cortical regions involved in decision making and executive function (e.g. ventromedial prefrontal cortex, dorsolateral prefrontal cortex) [19–27]. In particular, altered rsFC between the dorsal and ventral mesial prefrontal cortex (d/vMPFC), anterior cingulate cortex (ACC) and ventral striatum (VS) is most consistently observed across disorders of addiction such as cocaine [28], heroin [29], nicotine [30–33], and even internet addiction [32, 34, 35], but also in obsessive-compulsive disorder (OCD) [34]. Furthermore, vMPFC activity seems to be tightly linked to dMPFC activity [36, 37]. Thus, understanding whether and how deep rTMS targeting the MPFC influences the connected networks is critical to its potential clinical efficacy.

Study 1, we first assess rsFC between MPFC and striatum in a relatively large sample of healthy controls. In Study 2, we then ask whether inhibitory deep wide-field stimulation with an H7-coil positioned over the MPFC (which, given the non-focal nature of the H7-coil [38, 39], we have defined here as supplementary motor area (SMA), pre-SMA, and dMPFC) influences rsFC with VS in a separate group of healthy controls. We focused on VS given its aberrant rsFC observed in pathological disorders as well as in our findings in Study 1 of negative connectivity of MPFC with VS and positive connectivity of mid-cingulate with VS. We hypothesize that low-frequency inhibitory rTMS will decrease rsFC of the MPFC with VS.

## II. Methods

### A. Protocol Design and Participants

In Study 1, seed to whole brain intrinsic rsFC was examined for the mesial PFC (SMA, preSMA and dMPFC) and the mid-cingulate. For intrinsic baseline mapping, blood-oxygenation level dependent (BOLD) fMRI data was collected during rest (10 minutes, eyes open, watching white fixation cross on black screen) from 154 healthy volunteers (71 females; age 31±13 years) at the Wolfson Brain Imaging Centre, University of Cambridge, UK, with a Siemens Tim Trio 3T scanner and 32-channel head coil.

In Study 2, we used inhibitory, 1 Hz rTMS deep wide-field stimulation with an H7-coil targeting the mesial PFC. In order to examine the effects of rTMS on neural fluctuations, we used both ROI-to-ROI analyses and confirmed findings with independent component analysis (ICA). Resting state fMRI data (10 minutes, eyes open, watching white fixation cross) was collected immediately before and after rTMS (average time between rTMS end and fMRI start = 277 ± 27 seconds) in a separate group of 20 healthy volunteers (15 females; age 36 ± 12 years) at the National Institutes of Health (Bethesda, MD, USA) core fMRI Facility, with a Siemens Skyra 3T scanner and 32-channel head coil.

All subjects provided informed written consent. This study was approved by the Research Ethics Committee of the University of Cambridge and the Institutional Review Board of the National Institutes of Health.

### B. Transcranial Magnetic Stimulation with the H-coil (Study 2)

To modulate the excitability of deep frontal areas in Study 2, we used a Hesed coil type 7 (H7-coil). Its design aims at stimulating frontal brain regions (i.e., the PFC) and reaching deep brain regions without increasing the electric field levels in the more superficial cortical regions [6,40]. Deep TMS using other coils (e.g. classical double-cone coil) can be uncomfortable due to excessive stimulation of superficial structures and painful muscular contractions. The frames of the inner rim of H7-coil are also flexible to accommodate a variety of human skull shapes and allow a comfortable and closer fit of the coils to the scalp (Supplementary Figure S1).

We first found the hotspot and determined the active motor threshold (AMT) of the *tibialis anterior* muscle, as an area situated medially at a depth similar to our regions of interest (Figure 1A). The AMT was defined as the lowest intensity able to evoke a motor potential with an amplitude at least 200 µV above the background EMG activity of a 10% maximal voluntary contraction of the Tibialis anterior in 5 out of 10 consecutive trials. Repetitive TMS was delivered with a biphasic magnetic stimulator (Magstim Rapid2; The Magstim Company, Whitland, South West Wales, UK) with a frequency of 1 Hz and at 110% AMT intensity. Nine hundred pulses were administered over the MPFC, 5 cm anterior to the *tibialis anterior* hot-spot, for 15 min. When administered in accordance with current international guidelines, transcranial magnetic stimulation has been shown to be safe [41, 42], with few mild adverse effects, although we acknowledge that these safety guidelines are derived primarily from studies using conventional figure-8 coils.

**Fig. 1:**
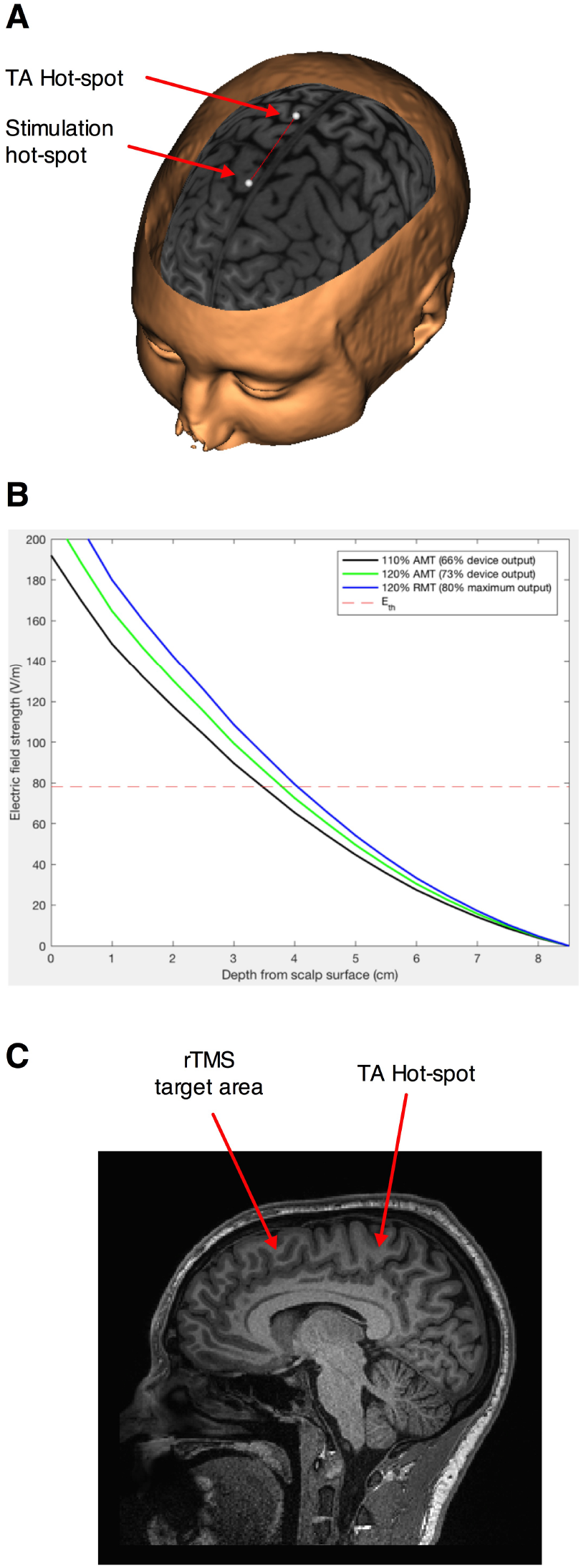
Stimulation paradigm. (A) Schematic representation of the movement of the projection of the geometric center of the H7 coil 5s cm in front of the empirically found hotspot for the left Tibialis anterior muscle. (B) Estimation of the induced electric field intensity with distance from the coil for stimulation at 110% of the active motor threshold (AMT)—our intensity of choice, and 120% AMT and 110% resting motor threshold—higher intensities distribution modeled for comparison. The dotted line represents the theoretical intensity of the induced electrical field for AMT. (C) Sagittal section showing the area in the dorso-mesial prefrontal cortex found at an equivalent depth to the Tibialis anterior motor representation.

We used medium intensity stimulation (i.e., 110% of the active motor threshold; average effective intensity 66 ± 8% of the maximum stimulator output) of the H7-coil, which would have penetrated effectively up to a depth of 3.5 cm from the surface of the scalp (Figure 1B), corresponding to the mesial PFC region (Figure 1C).

### C. Resting State Functional MRI

The following describes the resting state acquisitions and analyses used for Study 1 and 2. Acquisition Study 1: Functional images were acquired with a multi-echo echo planar imaging sequence with online reconstruction (repetition time (TR), 2.47 s; flip angle, 78°; matrix size 64×64; resolution 3.0×3.0×3.0 mm; FOV, 240 mm; 32 oblique slices, alternating slice acquisition slice thickness 3.75 mm with 10% gap; iPAT factor, 3; bandwidth (BW) = 1698 Hz/pixel; echo time (TE) = 12, 28, 44, and 60 ˙ms).

Study 2: Functional images were acquired with a multi-echo echo planar imaging sequence (TR, 2.47 s; flip angle, 70°; matrix size 70 × 60; in-plane resolution, 3.0 mm; FOV, 210 mm; 34 oblique slices, alternating slice acquisition slice thickness 3.0 mm with 0% gap; iPAT factor, 3; bandwidth (BW) = 2552 Hz/pixel; TE = 12, 28, 44, and 60 ms).

For both studies, anatomical images were acquired using a T1-weighted magnetization prepared rapid gradient echo (MPRAGE) sequence (76 × 240 field of view (FOV); resolution 1.0 × 1.0 × 1.0 mm; inversion time, 1100 ms).

### D. Preprocessing

The following processing and analyses apply to both resting state fMRI data unless stated otherwise. To enhance signal-to-noise ratio, we used multi-echo EPI sequence and independent component analysis (ICA), which allows data to be denoised for motion, physiological, and scanner artifacts in a robust manner based on physical principles [43]. Multi-echo independent component analysis (ME-ICA v2.5 beta6; http://afni.nimh.nih.gov) was used for data analysis and denoising. ME-ICA decomposes the functional data into independent components using FastICA. BOLD percent signal changes are linearly dependent on echo time (TE), a characteristic of the T2* decay. TE dependence of BOLD signal is measured using the pseudo-*F*–statistic, *κ*, with components that scale strongly with TE having high *κ* scores [44]. Non-BOLD components are TE independent and measured by the pseudo-*F*–statistic, *ρ*. Components are thus categorized as BOLD or non-BOLD based on their *κ* and *ρ* weightings, respectively. Non-BOLD components are removed by projection, robustly denoising data. Each individual’s denoised echo planar images were coregistered to their MPRAGE and normalized to the Montreal Neurological Institute (MNI) template. Spatial smoothing of the functional data was performed with a Gaussian kernel (full width half-maximum = 6 mm).

### E. Region of Interest (ROI)-Driven Analysis

We performed ROI-driven functional connectivity analysis using CONN-fMRI Functional Connectivity toolbox [45] for Statistical Parametric Mapping SPM8 (http://www.fil.ion.ucl.ac.uk>spm/software/spm8/), using denoised, coregistered, smoothed functional data. The time course for each voxel was temporally band-pass filtered (0.008 < *f* < 0.09 Hz). Each individual’s anatomical scan was segmented into grey matter, white matter and cerebrospinal fluid. Significant principle components of the signals from white matter and cerebrospinal fluid were removed.

#### 1) Study 1: Intrinsic Functional Connectivity Mapping

For intrinsic rsFC mapping in 154 healthy volunteers, ROI-to-whole brain connectivity was computed for mesial PFC and mid cingulate ROI’s. Connectivity maps were thresholded at FWE *p* < 0.05 whole brain corrected. Both positive and negative functional connectivity was examined across the whole brain. Anatomically-defined ROIs were manually created or altered using MarsBaR ROI toolbox [46] for SPM (see Section V-A for seed definitions)

#### 2) Study 2: Effects of rTMS: ROI-Based

To address the *a priori* hypothesis, ROI-to-ROI functional connectivity was first computed using Pearson’s correlation between BOLD time courses for mesial PFC with ventral striatum, both pre-and post-TMS. These were entered into a paired samples *t*-test to compare between pre-and post-TMS. For ROI-to-ROI functional connectivity analysis, *p* < 0.05 was considered significant. On an exploratory basis, to assess the impact of rTMS on rsFC of deeper structures such as the mid-cingulate which lies immediately below the mesial PFC, ROI-to-ROI functional connectivity of mesial PFC to mid cingulate and mid cingulate to VS were examined pre-and post-TMS. *p* < 0.025 was considered significant (Bonferonni corrected for multiple comparisons). The VS anatomical ROI has previously been used [47] and hand drawn using MRIcro (http://www.cabiatl.com/mricro/mricro/) based on a published definition of VS [48].

### F. Effects of rTMS: Independent Component Analysis (Study 2)

To confirm the ROI-to-ROI findings, we then conducted ICA. While ICA has been shown to engender statistically similar results as seed based approaches in healthy volunteers [49], ICA is a multivariate data-driven approach that requires fewer a priori assumptions and takes into account interacting networks. Therefore, if TMS affects larger scale neural networks, ICA should succeed in highlighting this. Denoised, coregistered, and smoothed functional data was entered into ICA analysis using FSL MELODIC 3.14 software (FMRIB, University of Oxford, UK; http://www.fmrib.ox.ac.uk/fsl/melodic2/index.html) that performs probabilistic ICA to decompose data into independently distributed spatial maps and associated time courses to identify independent component variables [50]. A high model order of 40 was used as a fair compromise between under-and over-fitting [51]. Multisession temporal concatenation was used to allow computation of unique temporal responses per subject/session. Comparisons between pre-and post-TMS was performed using FSL dual regression for reliable and robust [52] voxel-wise comparisons using nonparametric permutation testing with 5000 permutations and using threshold free cluster enhancement (TFCE) controlling for multiple comparisons [53]. Group differences of components that include MPFC were calculated with *p* < 0.05 thresholds.

## III. Results

### A. Baseline Mapping

Intrinsic resting state whole brain connectivity maps for mesial PFC and mid cingulate are displayed in Figure 2 and reported in Supplementary Table S1 and S2. Both positive and negative functional connectivity are displayed. Mesial PFC and mid cingulate showed opposite patterns of connectivity with ventral striatum: mesial PFC had negative but mid cingulate had positive functional connectivity with VS.

**Fig. 2:**
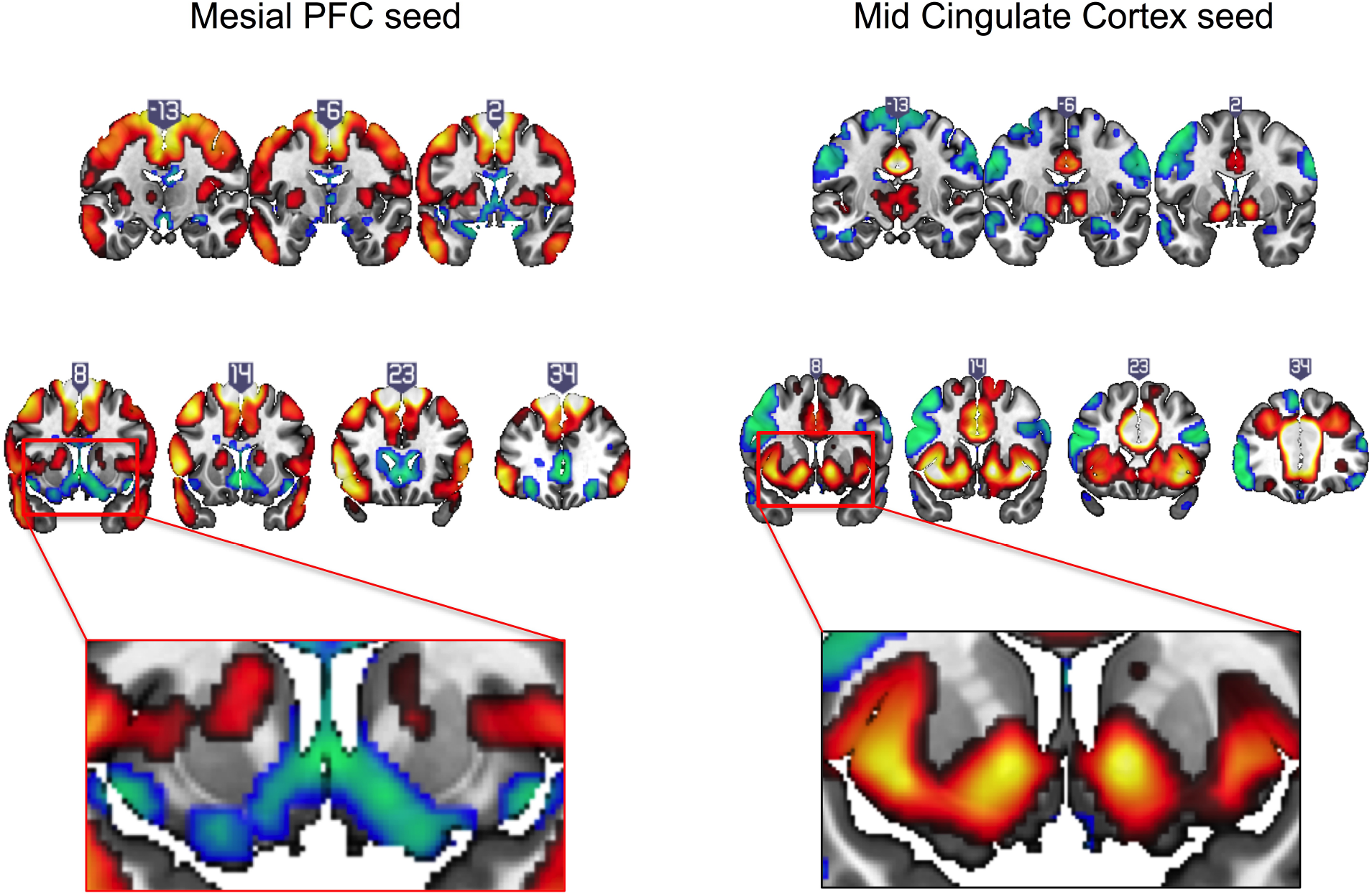
Intrinsic resting state connectivity maps for mesial prefrontal cortex (PFC) and mid cingulate cortex seeds to whole brain in healthy controls. Positive (yellow-red) and negative (green-blue) functional connectivity are displayed. The rectangular insets at *y* = 8 highlighting differences in direction of connectivity of the striatum are shown for the mesial PFC (bottom row, left) and mid cingulate (bottom row, right). Coronal images (*y*-values shown above image) are thresholded at whole brain family wise error corrected *p* < 0.05 on a standard MNI template.

### B. Effects of TMS

Focusing on our *a priori* hypothesis, we show that after rTMS, mesial PFC had reduced functional connectivity with ventral striatum (*t* = 2.201, *p* = 0.043) (Figure 3). We then show an effect on mid-cingulate functional connectivity with reduced functional connectivity following rTMS between the mesial PFC and mid-cingulate (*t* = 4.325, *p* = 0.001) and enhanced functional connectivity between mid-cingulate and VS (*t* = −2.495, *p* = 0.024).

**Fig. 3:**
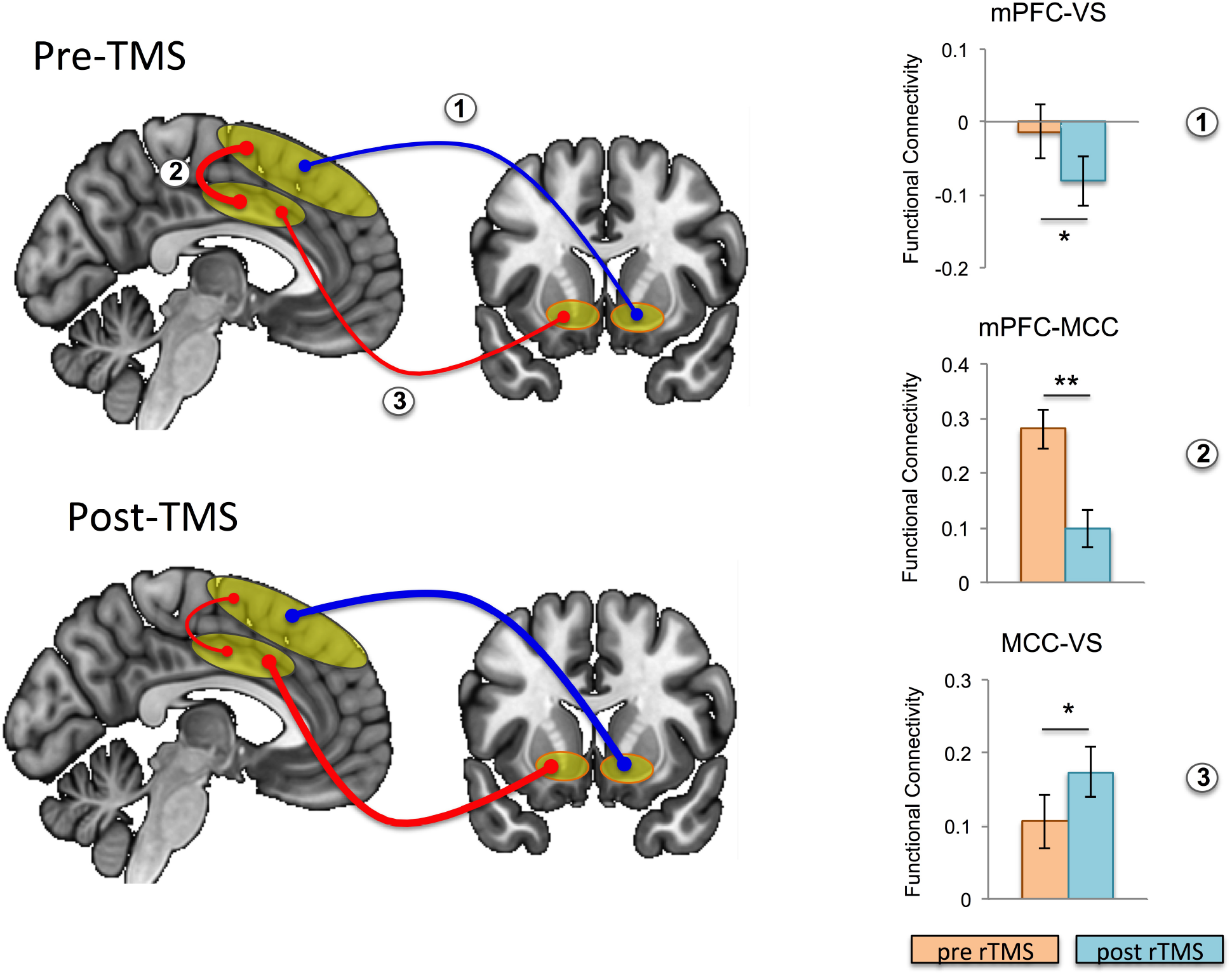
Effects of repetitive transcranial magnetic stimulation (rTMS) on intrinsic functional connectivity in healthy controls. Functional connectivity is schematically illustrated at baseline (i.e. pre-rTMS; top left) and post-rTMS (bottom left); pre-and post-rTMS effects on seed-to-seed functional connectivity are shown in the bar graphs. After rTMS, functional connectivity between mesial prefrontal cortex (mPFC) and ventral striatum (VS), and between mPFC and mid cingulate cortex (MCC) was reduced, while functional connectivity between MCC and VS was increased (the thickness of the arrows correspond to strength, and color to direction: red— positive connectivity, blue—negative connectivity). Error bars are shown as standard error of the mean. \**p* < 0.05, \*\**p* = 0.001

We conducted ICA on the resting state data pre-and post-rTMS to confirm our *a priori* hypothesis and analysis. Out of 40 components, three included prominent mesial frontal cortex (Figure 4). Of the three mesial frontal network components, dual regression revealed that one of these components (IC11) was significantly decreased post-rTMS (TFCE *p* = 0.036). The network included pre-SMA and SMA, dorsomedial PFC/dorsal cingulate, bilateral inferior frontal cortices, ventral caudate/ventral striatum, and midbrain.

**Fig. 4:**
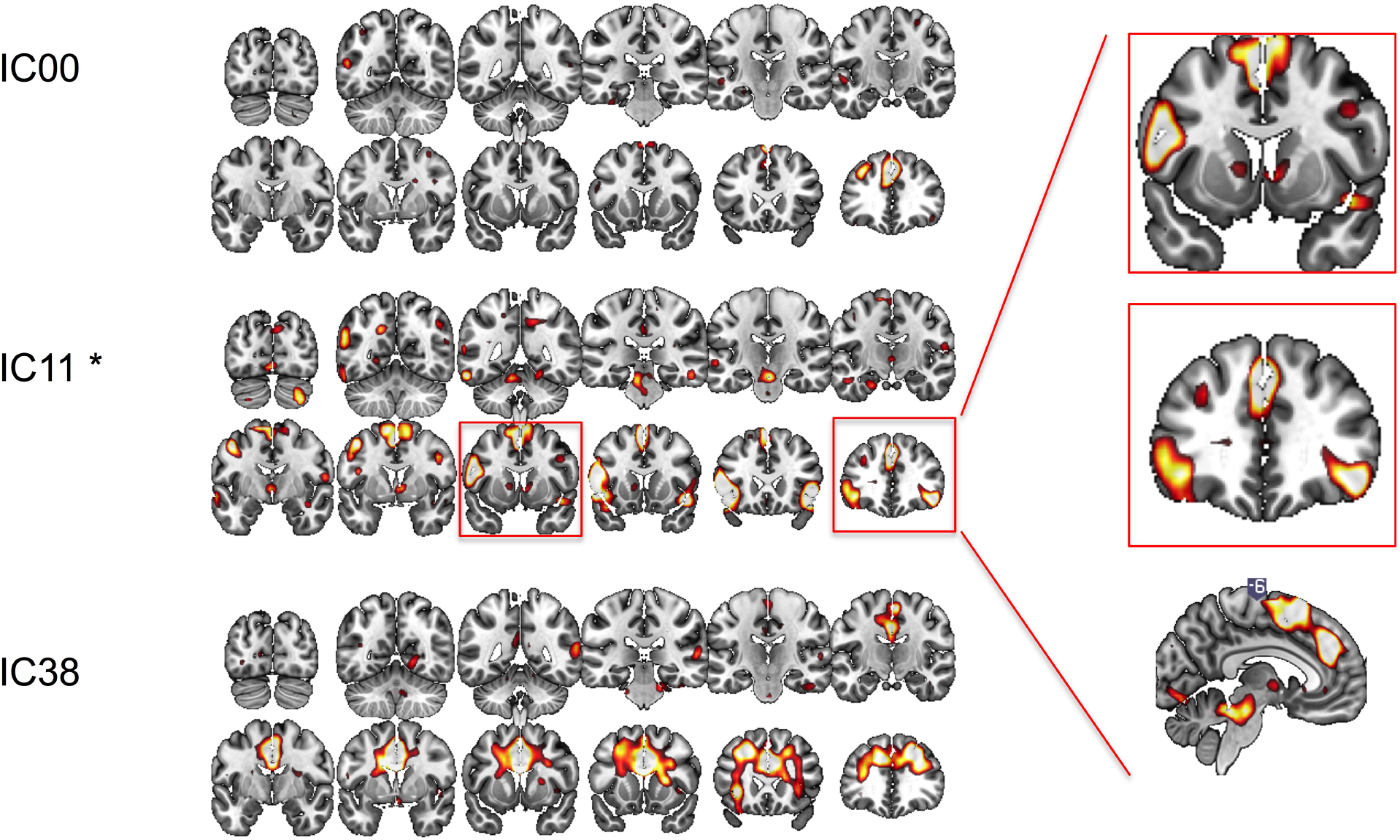
Functional connectivity at rest between different regions of interest explored with independent component analysis pre-and post-rTMS. Three components included prominent mesial-frontal cortex (IC00, IC11 and IC38). The insert shows IC11, which included supplementary motor area (SMA), pre-SMA, dorsomedial prefrontal cortex/dorsal cingulate, and ventral caudate/striatum, and bilateral inferior frontal cortices was significantly decreased post-rTMS. \**p* < 0.05

## IV. Discussion

We characterized the effects of deep wide-field mesial prefrontal rTMS on the resting-state functional network in healthy individuals. We first mapped intrinsic functional connectivity of mesial prefrontal and mid-cingulate cortical regions in a large sample of healthy volunteers. We found that intrinsic functional connectivity of the mesial PFC region of interest with ventral striatum was negative, whereas the intrinsic functional connectivity of mid-cingulate connectivity with ventral striatum was positive. Then, we show that deep wide-field inhibitory rTMS targeting the mesial PFC decreases rsFC between this broad mesial PFC region and the ventral striatum. These findings were further confirmed with ICA analysis, a data-driven approach. Based on the modeling of the magnetic field distribution, induced-electrical field decay, and the depth of the target region stimulated, we likely also inhibited directly the dorsal posterior regions of Brodmann Area 32, corresponding to dorsal anterior cingulate—a fact subsequently confirmed by the ICA analysis. Inhibitory rTMS also decreased functional connectivity of the “stopping” network including pre-SMA, right inferior frontal cortex, and ventral caudate. This is in line with previous reports, in which inhibitory rTMS (including continuous theta burst stimulation) targeting the pre-SMA with standard figure-of-eight coil has been shown to enhance motor response inhibition [54].

We also found effects of deep rTMS on connectivity between deeper structures such as the midcingulate cortex, which was unlikely to be directly stimulated with our stimulation parameters: decreased rsFC between the broad mesial PFC and mid-cingulate cortex, and, unexpectedly, enhanced rsFC between mid-cingulate cortex and ventral striatum. These findings suggest that while deep wide-field mesial prefrontal inhibitory rTMS might directly decrease the functional connectivity between the stimulated and the connected structures, the decreased influence from superficial cortical regions might indirectly enhance the intrinsic connectivity between remote structures (i.e., the mid-cingulate cortex and ventral striatum).

Application of rTMS to superficial cortical regions with the strongest negative functional connectivity with subgenual ACC has already been shown to be most clinically efficacious in reducing depression [55]. Thus, based on the deep cortical or subcortical structure of interest for a given disorder, appropriate superficial sites for rTMS can be selected based on intrinsic functional connectivity strengths and patterns. Since we demonstrate in our second study that there is an exaggeration of intrinsic functional connectivity strengths with deep inhibitory rTMS, detailed mapping of baseline connectivity patterns will inform the selection of rTMS targets with the aim to “normalize” aberrant underlying functional connectivity in disease states. The outcome of this modulation could be of interest in the treatment of disorders relying on the mesioprefrontal–cingulo–striatal circuit.

The H-coil series was originally designed to have a significant impact on deep structures, like the anterior cingulate cortex [6, 7]. It has been used with different degrees of success to treat depression [56, 57], alcohol use disorders [58], nicotine addiction [59], and even as adjunctive therapy in Parkinson’s disease [60], blepharospasm [61], and chronic migraine [62].

Due to the quick drop in TMS efficacy with increasing target depth [63], it has been proposed that any stimulation outside the primary motor cortex should be referenced to motor cortex excitability and adjusted to the target depth [64, 65]. The original assertion that the H-coil can modulate the activity of deep structures has been based mainly on calculating the intensity of the induced electrical field at different depths for a given stimulation intensity [40]. However, other factors can significantly influence the efficacy of rTMS, including the orientation of the coil [66–68] and the configuration of the subjacent and/or target cortex [69–73], as well as the secondary electrical fields generated at the boundary between the cerebrospinal fluid and the gray matter [74]. Subsequent studies of the distribution of the magnetic field generated by the H-coil revealed that the largest field intensity variation and hence, the functional effect covers first the mesial neuronal structures in close proximity to the coil, i.e., superior MF areas, like dMPFC, pre-SMA, SMA [40, 75–77], and only secondarily deeper structures such as the cingulate cortex if stimulation intensity is high enough [7, 40]. In order to reach the stimulation threshold of neurons, a total field of 30–100 V/m is needed, depending on the neurons [78]. Since focal coils, like flat 8-shaped or double-cone coils, produce very strong fields that decay fast as a function of distance, 500 V/m would be induced at 1 cm depth (i.e. scalp) for 50 V/m at 5 cm, which would be very uncomfortable due to superficial muscle contraction under the stimulated site [6]. According to our simulations (Figure 1B) using a spherical head model, the structure of the H7-coil induces only 150 V/m at 1cm in the same conditions, albeit at the cost of focality, making it more tolerable. In this study, we used medium intensity stimulation (i.e., 110% of the active motor threshold; average effective intensity 66 ± 8% of the maximum stimulator output), which would have stimulated a region of interest corresponding to the mesial PFC. This allowed us to influence directly the output of these areas and indirectly the activity of functionally linked structures [79–84]. Based on the simulated model of the target and depth reached using our stimulation parameters, we likely directly stimulated down to dorsal posterior regions of Brodmann Area 32 corresponding to dorsal anterior cingulate. However, it is unlikely that we directly stimulated the mid-cingulate; thus any change in connectivity observed in the mid-cingulate would likely be an indirect effect via changing the functional output of connected areas. Here, we extend the understanding of the effects of magnetic stimulation over the frontal lobe, following previous TMS studies investigating more superficial stimulation of the lateral frontal areas [55, 85–87]. Subsequent studies are indicated to investigate the influence of higher intensities and higher frequencies [88] on rsFC of frontal superficial and deep structures, when applied with coils designed to reach broader regions.

We delivered magnetic pulses at 1 Hz for 15 minutes. This frequency can induce a long term depression (LTD)-like effect in the targeted neuronal networks that outlasts the stimulation for a sufficient duration to assess the influence on resting-state fMRI [89–92]. By using low stimulation intensities, we effectively depressed the excitability of the superior mesial prefrontal areas and possibly also the dorsal posterior region of Brodmann Area 32 corresponding to dorsal anterior cingulate cortex. An LTD-like effect would thus decrease neuronal excitability in the mesial PFC, rendering it less responsive to incoming information. Decreased responsiveness would functionally decouple this region from both neighboring and deeper structures. Indeed, we found reduced functional connectivity of the broad mesial PFC with mid-cingulate, and between the broad mesial PFC and ventral striatum, with ICA confirming decreases in the network including mesial PFC, dorsal anterior cingulate and ventral caudate/ventral striatum. Since the fronto– striatal network relies on a dynamic equilibrium between its different parts [11, 93, 94], functionally “nudging” one part should entrain a reconfiguration of all functional connections, including functional connectivity between remote regions receiving projections from the stimulated region. This seems to be the case in our study: we found increased functional connectivity between the mid-cingulate area and ventral striatum after inhibiting the mesial PFC.

The outcome of this modulation could be of interest in treatment of disorders relying on the mesioprefrontal-cingulo-striatal circuit. In healthy humans, this circuit is involved in cognitive and emotional control, error and conflict monitoring [95–97], response inhibition [98], and positive and negative prediction error and anticipation [99–101]. Abnormal cortico–ventro striatal hyperconnectivity has been OCD [102–104] and addictions (for a review see [105]). In disorders of addiction, decreased functional connectivity between the ventral striatum and the cingulate cortex bilaterally is commonly observed [29, 32], with enhanced dorsal cingulate and ventral striatal activity in the context of drug cues [106]. Numerous targets had been proposed for invasive deep brain stimulation aimed at correcting these imbalances, including the anterior limb of the internal capsule [107], subthalamic nucleus [108], and ventral striatum/nucleus accumbens [109]. In order to avoid the risks of an invasive procedure, studies have explored stimulating other nodes of these networks that are accessible to TMS at the surface of the brain. Stimulation of the dorsolateral prefrontal cortex, is (arguably [56, 57]) successful in treatment-resistant major depressive disorder [4, 110], with modest results in OCD [111]. On the other hand, stimulation of the dorso-medial prefrontal cortex [112] or pre-SMA/SMA complex [113–115] seems slightly more encouraging. Notably, there is no gold standard yet for the frequencies to be used. The stimulation frequencies used thus far in most studies cover a wide range including continuous delivery at 1 Hz, or intermittently at 10 or 18 Hz in 5 s trains separated by breaks of 10 s. While 1 Hz stimulation is known to induce LTD-like effects, the mechanism of action and the eventual outcome of other multiple medium-frequency trains is still open to debate and investigation [116, 117].

Wide inhibitory stimulation of the dorso-mesial areas of the frontal lobe might have both clinical and mechanistic benefit. Wider superficial stimulation has a clear clinical benefit allowing a reduction in the intensity of the stimulation with deeper stimulation, thus increasing patients’ comfort and adherence by decreasing superficial muscle contraction, and minimizing risks. Aberrant activity in networks in psychiatric disorders may affect broader regions that can be targeted via wide inhibitory stimulation. We show that stimulation that is both wide and deep is associated with decreased connectivity between the mesial prefrontal areas and deeper structures (like the mid-cingulate areas and ventral striatum), with possibly a secondary effect of increasing connectivity between cingulate and ventral striatum. Wider stimulation will also have a broader effect on multiple neural regions, impacting a wide range of cognitive functions. Using the H7-coil with inhibitory rTMS is thus consistent with both inhibition of the pre-SMA shown to enhance motor response inhibition [54] and decreased dorsal cingulate activity associated with drug cue reactivity [106]. Therefore, the H7-coil has the capacity to both enhance the response inhibition associated with the stopping network in disorders of addiction, and decrease drug cue reactivity associated with the dorsal cingulate and ventral striatum. However, it is unclear whether decreasing dorsal cingulate activity across all conditions would be the optimal approach, as resting state functional connectivity between cingulate and ventral striatal regions are commonly decreased in disorders of addiction. Further studies investigating a state-specific effect of rTMS may be relevant with pairing H-coil stimulation with drug cues with or without concurrent response inhibition. It also remains to be established whether our findings are specific to wide-field deep rTMS or whether focal deep rTMS (which is be more difficult to tolerate) would show similar rsFC pattern changes within cingulate regions.

This study is not without limitations. We tested subjects pre-and post-real rTMS always in the same order. While we did not have a sham control, we note that our findings revealed both increases and decreases in connectivity—suggesting that an order effect is unlikely to account for these observations. The localization of the peak stimulus effect is also more difficult with the H-coil, since the coils’ positions inside the helmet do not generate a magnetic field flux with a well-localized maximum. Subsequent studies testing higher frequencies and/or intensities are indicated, as well as repeated stimulation sessions (over minimum 4 weeks) in preparation for clinical trials.

We highlight that non-invasive wide and deep inhibitory brain stimulation appears to decrease the underlying functional connectivity of regions immediately within the stimulation zone while enhancing functional connectivity of deeper structures such as midcingulate to ventral striatum. This unexpected finding might be related to the decreased influence from superficial cortical regions via decreased cortico-cortical connectivity. A deep wide-field coil allows both greater tolerability and the capacity to influence multiple relevant neural regions and cognitive functions. These dissociable findings may be relevant particularly to disorders of addiction and OCD, and have implications for designing interventional deep rTMS studies.

## V. Supplementary Methods

### A. Seed Definitions

The anatomically defined ROIs which were manually created or altered using MarsBaR ROI toolbox [46] for SPM. The broad mesial prefrontal ROI was defined with the posterior border as the extent of the SMA and the anterior border as the anterior extent of the dorsal ACC. The statistics for positive and negative functional connectivity of the medial prefrontal cortex seed are reported in Supplementary Table S1.

**Table S1:** Statistics for positive and negative functional connectivity of medial prefrontal cortex (MPFC) seed.

| | $p(\text{FWE-corr})$ | $K$ | $T$ | $Z$ | $x$ | $y$ | $z$ |
| --- | --- | --- | --- | --- | --- | --- | --- |
| **MPFC positive** |  |  |  |  |  |  |  |
| MPFC | < 0.001 | 36108 | 36.7 | > 8 | -10 | 19 | 61 |
|  |  |  | 35.73 | > 8 | -10 | 31 | 56 |
|  |  |  | 33.56 | > 8 | -1 | 7 | 70 |
| Cerebellum | < 0.001 | 2608 | 16.49 | > 8 | 29 | -79 | -32 |
|  |  |  | 14.27 | > 8 | 36 | -62 | -28 |
|  |  |  | 11.2 | > 8 | 34 | -60 | -53 |
| Cerebellum | < 0.001 | 1245 | 12.24 | > 8 | -27 | -74 | -30 |
|  |  |  | 10 | > 8 | -34 | -65 | -25 |
|  |  |  | 5.52 | 5.26 | -24 | -58 | -23 |
| Lateral Parietal | < 0.001 | 625 | 11.28 | > 8 | 57 | -55 | 35 |
| Thalamus | < 0.001 | 137 | 9.89 | > 8 | -8 | -20 | 3 |
| Cerebellum | < 0.001 | 24 | 7.44 | 6.86 | 6 | -55 | -42 |
|  |  |  | 5.87 | 5.57 | 6 | -60 | -49 |
| Midbrain | < 0.001 | 34 | 7.02 | 6.53 | -1 | -23 | -30 |
| Cerebellum | < 0.001 | 25 | 6.95 | 6.47 | -8 | -62 | -42 |
| Cerebellum | < 0.001 | 46 | 6.6 | 6.18 | -1 | -53 | -9 |
| Cerebellum | < 0.001 | 103 | 6.58 | 6.17 | -34 | -62 | -56 |
|  |  |  | 5.95 | 5.63 | -41 | -55 | -51 |
| Thalamus | < 0.001 | 24 | 6.33 | 5.95 | 10 | -18 | 3 |
| Medial Parietal | < 0.001 | 24 | 5.58 | 5.32 | -10 | -51 | 33 |
| Cerebellum | 0.011 | 4 | 5.4 | 5.15 | -24 | -39 | -28 |
| **MPFC negative** |  |  |  |  |  |  |  |
| Medial Parietal | < 0.001 | 13212 | 14.78 | > 8 | -17 | -60 | 21 |
|  |  |  | 14.47 | > 8 | 15 | -51 | 19 |
|  |  |  | 13.75 | > 8 | 15 | −67 | 28 |
| Lateral OFC | < 0.001 | 136 | 9.66 | > 8 | −27 | 35 | −14 |
| Lateral OFC | < 0.001 | 106 | 9.52 | > 8 | 22 | 33 | −16 |
| Cerebellum | < 0.001 | 237 | 8.95 | > 8 | 1 | −46 | −30 |
|  |  |  | 8.25 | 7.49 | −10 | −48 | −46 |
| Temporal | < 0.001 | 150 | 8.23 | 7.48 | −50 | −60 | −2 |
| Cerebellum | < 0.001 | 102 | 7.79 | 7.14 | 13 | −44 | −49 |
| Temporal | 0.001 | 18 | 6 | 5.68 | 48 | −51 | −4 |
| Cerebellum | 0.001 | 17 | 5.8 | 5.5 | −6 | −83 | −37 |
| Lateral Prefrontal | 0.001 | 19 | 5.49 | 5.24 | 41 | 33 | 19 |

Abbreviations: $p$ (FWE-corr), whole brain ( $p < 0.05$ ) family-wise error corrected $p$ -value; $K$ , cluster size; $T$ , $T$ -statistic; $Z$ , $Z$ -score; $xyz$ , peak voxel coordinates; OFC, orbitofrontal cortex.

The mid cingulate ROI had an anterior border of the posterior end of the genu of the corpus callosum and posterior border a vertical line through the anterior commissure—the same as the pre-SMA posterior border. The statistics for positive and negative functional connectivity of the mid-cingulate cortex seed are reported in Supplementary Table S2.

**Table S2:** Statistics for positive and negative functional connectivity of mid-cingulate cortex seed.

| | $p$(FWE-corr) | $K$ | $T$ | $Z$ | $x$ | $y$ | $z$ |
| --- | --- | --- | --- | --- | --- | --- | --- |
| **Mid Cingulate positive** |  |  |  |  |  |  |  |
| Mid Cingulate | < 0.001 | 30314 | 54.16 | > 8 | −6 | 14 | 35 |
|  |  |  | 52.52 | > 8 | −3 | 7 | 40 |
|  |  |  | 51.1 | > 8 | 3 | 17 | 35 |
| Dorsolateral PFC | < 0.001 | 513 | 13.04 | > 8 | 29 | 45 | 28 |
|  |  |  | 5.97 | 5.65 | 38 | 42 | 14 |
| Cerebellum | < 0.001 | 300 | 10.92 | > 8 | −31 | −53 | −51 |
|  |  |  | 6.14 | 5.8 | −17 | −62 | −53 |
| Temporal | < 0.001 | 68 | 10.53 | > 8 | −20 | −41 | 0 |
| Cerebellum | < 0.001 | 233 | 10.11 | > 8 | −34 | −53 | −28 |
|  |  |  | 6.94 | 6.46 | −43 | −58 | −30 |
| Temporal | < 0.001 | 57 | 10.02 | > 8 | 20 | −39 | 3 |
| Cerebellum | < 0.001 | 356 | 9.9 | > 8 | 34 | −51 | −51 |
|  |  |  | 7.61 | 6.99 | 20 | −48 | −56 |
|  |  |  | 7.05 | 6.55 | 13 | −60 | −51 |
| Cerebellum | < 0.001 | 183 | 9.52 | > 8 | 34 | −48 | −30 |
|  |  |  | 7.34 | 6.78 | 43 | −55 | −32 |
|  |  |  | 5.33 | 5.09 | 22 | −60 | −23 |
| Inferior Frontal | < 0.001 | 90 | 7.69 | 7.06 | −29 | 35 | −14 |
| Occipital | < 0.001 | 52 | 7.01 | 6.52 | −1 | −58 | −7 |
| Cerebellum | 0.001 | 20 | 6.42 | 6.03 | −3 | −34 | −42 |
| Occipital | 0.001 | 21 | 6.12 | 5.78 | −24 | −67 | 7 |
| Occipital | 0.011 | 4 | 5.26 | 5.04 | 52 | −60 | 3 |
| Occipital | 0.002 | 12 | 5.26 | 5.03 | 24 | −60 | 7 |
| **Mid Cingulate negative** |  |  |  |  |  |  |  |
| Cerebellum | < 0.001 | 12058 | 12.82 | > 8 | −13 | −74 | −30 |
|  |  |  | 11.61 | > 8 | −31 | −74 | −32 |
|  |  |  | 11.31 | > 8 | 41 | −62 | 59 |
| Temporal Cortex | < 0.001 | 707 | 11.5 | > 8 | 66 | −27 | −7 |
|  |  |  | 6.3 | 5.93 | 66 | −48 | −14 |
|  |  |  | 5.34 | 5.1 | 62 | −2 | −18 |
| Dorsolateral PFC | < 0.001 | 494 | 11.17 | > 8 | −43 | 10 | 56 |
|  |  |  | 6.79 | 6.34 | −57 | 21 | 21 |
| Dorsolateral PFC | < 0.001 | 1278 | 10.83 | > 8 | 48 | 14 | 52 |
|  |  |  | 10.42 | > 8 | 52 | 21 | 45 |
|  |  |  | 7.43 | 6.85 | 43 | 17 | 19 |
| Cerebellum | < 0.001 | 396 | 9.57 | > 8 | −6 | −55 | −37 |
|  |  |  | 8.61 | 7.76 | −6 | −58 | −49 |
|  |  |  | 8.61 | 7.75 | 1 | −58 | −44 |
| Mid Cingulate | < 0.001 | 301 | 8.31 | 7.53 | 3 | 12 | 12 |
|  |  |  | 8.25 | 7.49 | −6 | 14 | 12 |
|  |  |  | 7.9 | 7.22 | 6 | 3 | 17 |
| Temporal | < 0.001 | 342 | 7.95 | 7.26 | −64 | −30 | −4 |
|  |  |  | 6.9 | 6.43 | −66 | −39 | −2 |
| Frontal Polar | < 0.001 | 414 | 7.94 | 7.26 | 34 | 66 | 0 |
|  |  |  | 7.05 | 6.55 | 43 | 54 | −14 |
|  |  |  | 6.01 | 5.68 | 48 | 33 | −11 |
| Cerebellum | < 0.001 | 157 | 7.91 | 7.23 | 8 | −37 | −16 |
|  |  |  | 6.91 | 6.44 | 17 | −27 | −30 |
|  |  |  | 6.6 | 6.18 | 17 | −14 | −30 |
| Inferior Frontal | < 0.001 | 289 | 7.36 | 6.8 | −50 | 38 | −9 |
| Midbrain | < 0.001 | 171 | 7.35 | 6.79 | −6 | −7 | −21 |
|  |  |  | 5.18 | 4.96 | 1 | 5 | −21 |
| Thalamus | < 0.001 | 110 | 7.32 | 6.77 | 29 | −34 | 12 |
|  |  |  | 6.54 | 6.13 | 24 | −30 | 5 |
| Thalamus | < 0.001 | 92 | 6.99 | 6.5 | −27 | −32 | 5 |
|  |  |  | 6.47 | 6.08 | −36 | −32 | 0 |
|  |  |  | 6.34 | 5.96 | −29 | −39 | 12 |
| Dorsomedial PFC | < 0.001 | 111 | 6.93 | 6.45 | 1 | 40 | 45 |
| Cerebellum | < 0.001 | 81 | 6.49 | 6.09 | −13 | −32 | −18 |
|  |  |  | 6.21 | 5.85 | −6 | −44 | −7 |
|  |  |  | 5.42 | 5.17 | −17 | −27 | −32 |
| Temporal | < 0.001 | 93 | 6.22 | 5.86 | −59 | −11 | −21 |
|  | < 0.001 | 29 | 6.03 | 5.7 | −22 | −20 | 31 |
| Cerebellum | 0.002 | 14 | 6.01 | 5.69 | 22 | −34 | −42 |
| Frontal | 0.001 | 15 | 5.92 | 5.61 | −20 | 26 | 12 |
| Medial Parietal | < 0.001 | 27 | 5.82 | 5.52 | 3 | −74 | 54 |
| Posterior Cingulate | 0.003 | 11 | 5.79 | 5.5 | 1 | −51 | 7 |
| Frontal | 0.007 | 6 | 5.72 | 5.44 | −29 | 31 | 17 |
| Mid Cingulate | < 0.001 | 40 | 5.66 | 5.38 | 17 | −14 | 26 |
|  |  |  | 5.46 | 5.21 | 10 | −16 | 21 |
|  |  |  | 5.31 | 5.08 | 15 | −30 | 21 |

Abbreviations: $p$ (FWE-corr), whole brain ( $p < 0.05$ ) family-wise error corrected $p$ -value; $K$ , cluster size; $T$ , $T$ -statistic; $Z$ , $Z$ -score; $xyz$ , peak voxel coordinates; PFC, prefrontal cortex.

### B. Electric Field Induced by H7 Coil

The head, coils, and electric field were modeled with the electromagnetic finite element package MagNet (Infolytica, Inc., Canada). The electric field simulation methods were previously described in detail and validated experimentally [118]. Briefly, the human head was modeled as a homogeneous sphere with radius of 8.5 cm and isotropic conductivity of 0.33 S/m. The H7 coil consisted of two adjacent wings fixed at a relative angle of 90 degrees; each wings consisted of two layers of concentric elliptical windings with major axis ranging from 75–140 mm, and minor axis ranging from 70−−125 mm; each layer has 4 turns. The coil windings were modeled as stranded copper wires with cross-sectional diameter of 4 mm. The electric field distribution was computed using the MagNet Time Harmonic solver, and scaled to match the output of a Magstim Rapid2 device. We estimated the electric field threshold for neural activation, *E*_th_, using a linear model of neuronal response to TMS [118].

**Fig. S1:**
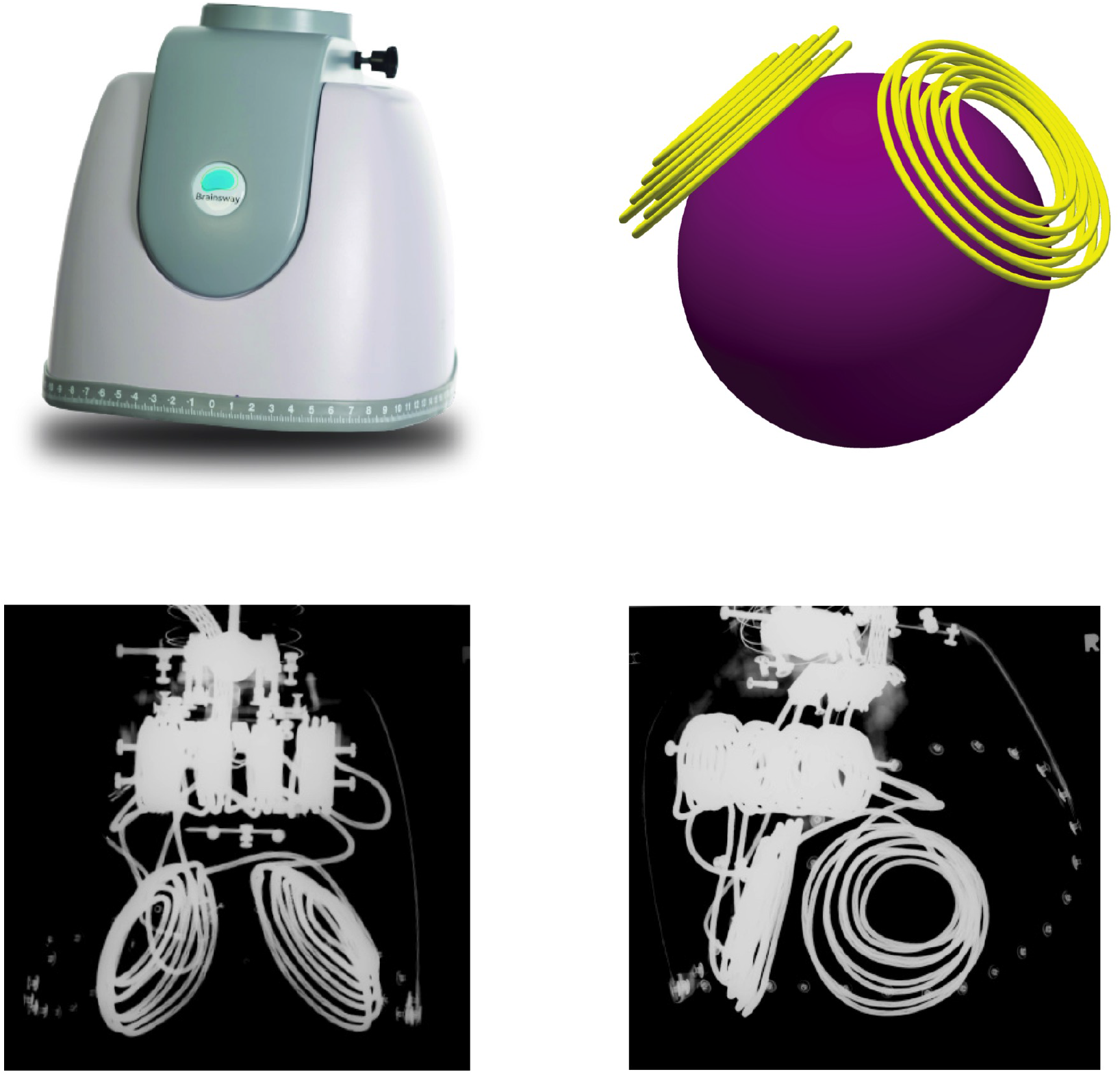
The H7 helmet with the internal wiring and the spherical head model used for calculating the induced-electrical field decay with distance from the coil.

## Acknowledgments

We thank our subjects for taking part in this study and the NMR Center personnel for the efficient assistance.

## References

[1] F. S. Bersani, A. Minichino, P. G. Enticott, L. Mazzarini, N. Khan, G. Antonacci, R. N. Raccah, M. Salviati, R. Delle Chiaie, G. Bersani, P. B. Fitzgerald, and M. Biondi, “Deep transcranial magnetic stimulation as a treatment for psychiatric disorders: a comprehensive review,” Eur Psychiatry, vol. 28, no. 1, pp. 30–39, 2013.

[2] Y. Roth, F. Padberg, and A. Zangen, Transcranial Brain Stimulation for Treatment of Psychiatric Disorders. Karger: Basel, 2007, ch. Transcranial Magnetic Stimulation of Deep Brain Regions: Principles and Methods, pp. 204–224.

[3] M. T. Berlim, A. McGirr, F. Van den Eynde, M. P. Fleck, and P. Giacobbe, “Effectiveness and acceptability of deep brain stimulation (DBS) of the subgenual cingulate cortex for treatment-resistant depression: a systematic review and exploratory meta-analysis,” J Affect Disord, vol. 159, pp. 31–38, 2014.

[4] M. T. Berlim, F. Van den Eynde, S. Tovar-Perdomo, E. Chachamovich, A. Zangen, and G. Turecki, “Augmenting antidepressants with deep transcranial magnetic stimulation (dTMS) in treatment-resistant major depression,” World J Biol Psychiatry, vol. 15, no. 7, pp. 570–578, 2014.

[5] J. P. Lefaucheur, N. André-Obadia, A. Antal, S. S. Ayache, C. Baeken, D. H. Benninger, R. M. Cantello, M. Cincotta, M. de Carvalho, D. De Ridder, H. Devanne, V. Di Lazzaro, S. R. Filipović, F. C. Hummel, S. K. Jääskeläinen, V. K. Kimiskidis, G. Koch, B. Langguth, T. Nyffeler, A. Oliviero, F. Padberg, E. Poulet, S. Rossi, P. M. Rossini, J. C. Rothwell, C. Schönfeldt-Lecuona, H. R. Siebner, C. W. Slotema, C. J. Stagg, J. Valls-Sole, U. Zie-mann, W. Paulus, and L. Garcia-Larrea, “Evidence-based guidelines on the therapeutic use of repetitive transcranial magnetic stimulation (rTMS),” Clin Neurophysiol, vol. 125, no. 11, pp. 2150–2206, 2014.

[6] Y. Roth, A. Zangen, and M. Hallett, “A coil design for transcranial magnetic stimulation of deep brain regions,” J Clin Neurophysiol, vol. 19, no. 4, pp. 361–370, 2002.

[7] A. Zangen, Y. Roth, B. Voller, and M. Hallett, “Transcranial magnetic stimulation of deep brain regions: evidence for efficacy of the H-coil,” Clin Neurophysiol, vol. 116, no. 4, pp. 775–779, 2005.

[8] E. K. Miller, “The prefrontal cortex and cognitive control,” Nat Rev Neurosci, vol. 1, no. 1, pp. 59–65, 2000.

[9] O. Hikosaka and M. Isoda, “Switching from automatic to controlled behavior: cortico-basal ganglia mechanisms,” Trends Cogn Sci, vol. 14, no. 4, pp. 154–161, 2010.

[10] I. Lee and C. H. Lee, “Contextual behavior and neural circuits,” Front Neural Circuits, vol. 7, no. 84, 2013.

[11] L. S. Morris, P. Kundu, N. Dowell, D. J. Mechelmans, P. Favre, M. A. Irvine, T. W. Robbins, N. Daw, E. T. Bullmore, N. A. Harrison, and V. Voon, “Fronto-striatal organization: Defining functional and microstructural substrates of behavioural flexibility,” Cortex, vol. 74, pp. 118–133, 2016.

[12] B. Biswal, F. Z. Yetkin, V. M. Haughton, and J. S. Hyde, “Functional connectivity in the motor cortex of resting human brain using echo-planar MRI,” Magn Reson Med, vol. 34, no. 4, pp. 537–541, 1995.

[13] B. B. Biswal, J. Van Kylen, and J. S. Hyde, “Simultaneous assessment of flow and BOLD signals in resting-state functional connectivity maps,” NMR Biomed, vol. 10, no. 4–5, pp. 165–170, 1997.

[14] J. L. Vincent, G. H. Patel, M. D. Fox, A. Z. Snyder, J. T. Baker, D. C. Van Essen, J. M. Zempel, L. H. Snyder, M. Corbetta, and M. E. Raichle, “Intrinsic functional architecture in the anaesthetized monkey brain,” Nature, vol. 447, no. 7140, pp. 83–86, 2007.

[15] S. M. Smith, P. T. Fox, K. L. Miller, D. C. Glahn, P. M. Fox, C. E. Mackay, N. Filippini, K. E. Watkins, R. Toro, A. R. Laird, and C. F. Beckmann, “Correspondence of the brain’s functional architecture during activation and rest,” Proc Natl Acad Sci U S A, vol. 106, no. 31, pp. 13 040–13 045, 2009.

[16] W. W. Seeley, V. Menon, A. F. Schatzberg, J. Keller, G. H. Glover, H. Kenna, A. L. Reiss, and M. D. Greicius, “Dissociable intrinsic connectivity networks for salience processing and executive control,” J Neurosci, vol. 27, no. 9, pp. 2349–2356, 2007.

[17] M. E. Raichle and M. A. Mintun, “Brain work and brain imaging,” Annu Rev Neurosci, vol. 29, pp. 449–476, 2006.

[18] M. Greicius, “Resting-state functional connectivity in neuropsychiatric disorders,” Curr Opin Neurol, vol. 21, no. 4, pp. 424–430, 2008.

[19] N. A. Fineberg, M. N. Potenza, S. R. Chamberlain, H. A. Berlin, L. Menzies, A. Bechara, B. J. Sahakian, T. W. Robbins, E. T. Bullmore, and E. Hollander, “Probing compulsive and impulsive behaviors, from animal models to endophenotypes: a narrative review,” Neuropsychopharmacology, vol. 35, no. 3, pp. 591–604, 2010.

[20] J. Feil, D. Sheppard, P. B. Fitzgerald, M. Yücel, D. I. Lubman, and J. L. Bradshaw, “Addiction, compulsive drug seeking, and the role of frontostriatal mechanisms in regulating inhibitory control,” Neurosci Biobehav Rev, vol. 35, no. 2, pp. 248–275, 2010.

[21] J. Camchong, A. W. MacDonald, 3rd, B. Nelson, C. Bell, B. A. Mueller, S. Specker, and K. O. Lim, “Frontal hyperconnectivity related to discounting and reversal learning in cocaine subjects,” Biol Psychiatry, vol. 69, no. 11, pp. 1117–1123, 2011.

[22] Y. Sakai, J. Narumoto, S. Nishida, T. Nakamae, K. Yamada, T. Nishimura, and K. Fukui, “Corticostriatal functional connectivity in non-medicated patients with obsessive-compulsive disorder,” Eur Psychiatry, vol. 26, no. 7, pp. 463–469, 2011.

[23] C. E. Wilcox, T. M. Teshiba, F. Merideth, J. Ling, and A. R. Mayer, “Enhanced cue reactivity and fronto-striatal functional connectivity in cocaine use disorders,” Drug Alcohol Depend, vol. 115, no. 1–2, pp. 137–144, 2011.

[24] A. B. Konova, S. J. Moeller, and R. Z. Goldstein, “Common and distinct neural targets of treatment: changing brain function in substance addiction,” Neurosci Biobehav Rev, vol. 37, no. 10 Pt 2, pp. 2806–2817, 2013.

[25] S. Koehler, S. Ovadia-Caro, E. van der Meer, A. Villringer, A. Heinz, N. Romanczuk-Seiferth, and D. S. Margulies, “Increased functional connectivity between prefrontal cortex and reward system in pathological gambling,” PLoS One, vol. 8, no. 12, e84565, 2013.

[26] D. Tomasi and N. D. Volkow, “Striatocortical pathway dysfunction in addiction and obesity: differences and similarities,” Crit Rev Biochem Mol Biol, vol. 48, no. 1, pp. 1–19, 2013.

[27] Y. Abe, Y. Sakai, S. Nishida, T. Nakamae, K. Yamada, K. Fukui, and J. Narumoto, “Hyper-influence of the orbitofrontal cortex over the ventral striatum in obsessive-compulsive disorder,” Eur Neuropsychopharmacol, vol. 25, no. 11, pp. 1898–1905, 2015.

[28] K. M. Wisner, E. H. Patzelt, K. O. Lim, and A. W. MacDonald, 3rd, “An intrinsic connectivity network approach to insula-derived dysfunctions among cocaine users,” Am J Drug Alcohol Abuse, vol. 39, no. 6, pp. 403–413, 2013.

[29] W. Wang, Y. R. Wang, W. Qin, K. Yuan, J. Tian, Q. Li, L. Y. Yang, L. Lu, and Y. M. Guo, “Changes in functional connectivity of ventral anterior cingulate cortex in heroin abusers,” Chin Med J (Engl), vol. 123, no. 12, pp. 1582–1588, 2010.

[30] L. E. Hong, C. A. Hodgkinson, Y. Yang, H. Sampath, T. J. Ross, B. Buchholz, B. J. Salmeron, V. Srivastava, G. K. Thaker, D. Goldman, and E. A. Stein, “A genetically modulated, intrinsic cingulate circuit supports human nicotine addiction,” Proc Natl Acad Sci U S A, vol. 107, no. 30, pp. 13 509–13 514, 2010.

[31] L. E. Hong, H. Gu, Y. Yang, T. J. Ross, B. J. Salmeron, B. Buchholz, G. K. Thaker, and E. A. Stein, “Association of nicotine addiction and nicotine’s actions with separate cingulate cortex functional circuits,” Arch Gen Psychiatry, vol. 66, no. 4, pp. 431–441, 2009.

[32] F. Lin, Y. Zhou, Y. Du, Z. Zhao, L. Qin, J. Xu, and H. Lei, “Aberrant corticostriatal functional circuits in adolescents with internet addiction disorder,” Front Hum Neurosci, vol. 9, no. 356, 2015.

[33] K. Yuan, D. Yu, Y. Bi, Y. Li, Y. Guan, J. Liu, Y. Zhang, W. Qin, X. Lu, and J. Tian, “The implication of frontostriatal circuits in young smokers: A resting-state study,” Hum Brain Mapp, vol. 37, no. 6, pp. 2013–2026, 2016.

[34] L. F. Fontenelle, B. J. Harrison, J. Pujol, C. G. Davey, A. Fornito, E. Bora, C. Pantelis, and M. Yücel, “Brain functional connectivity during induced sadness in patients with obsessive-compulsive disorder,” J Psychiatry Neurosci, vol. 37, no. 4, pp. 231–240, 2012.

[35] N. Ma, Y. Liu, N. Li, C. X. Wang, H. Zhang, X. F. Jiang, H. S. Xu, X. M. Fu, X. Hu, and D. R. Zhang, “Addiction related alteration in resting-state brain connectivity,” Neuroimage, vol. 49, no. 1, pp. 738–744, 2010.

[36] A. J. Jasinska, B. T. Chen, A. Bonci, and E. A. Stein, “Dorsal medial prefrontal cortex (MPFC) circuitry in rodent models of cocaine use: implications for drug addiction therapies,” Addict Biol, vol. 20, no. 2, pp. 215–226, 2015.

[37] A. J. Jasinska, E. A. Stein, J. Kaiser, M. J. Naumer, and Y. Yalachkov, “Factors modulating neural reactivity to drug cues in addiction: a survey of human neuroimaging studies,” Neurosci Biobehav Rev, vol. 38, pp. 1–16, 2014.

[38] V. Guadagnin, M. Parazzini, S. Fiocchi, I. Liorni, and P. Ravazzani, “Deep transcranial magnetic stimulation: modeling of different coil configurations,” IEEE Trans Biomed Eng, vol. 63, no. 7, pp. 1543–1550, 2016.

[39] S. Fiocchi, M. Longhi, P. Ravazzani, Y. Roth, A. Zangen, and M. Parazzini, “Modelling of the electric field distribution in deep transcranial magnetic stimulation in the adolescence, in the adulthood, and in the old age,” Comput Math Methods Med, vol. 2016, no. 9039613, 2016.

[40] Y. Roth, A. Amir, Y. Levkovitz, and A. Zangen, “Three-dimensional distribution of the electric field induced in the brain by transcranial magnetic stimulation using figure-8 and deep H-coils,” J Clin Neurophysiol, vol. 24, no. 1, pp. 31–38, 2007.

[41] Y. Levkovitz, Y. Roth, E. V. Harel, Y. Braw, A. Sheer, and A. Zangen, “A randomized controlled feasibility and safety study of deep transcranial magnetic stimulation,” Clin Neurophysiol, vol. 118, no. 12, pp. 2730–2744, 2007.

[42] S. Rossi, M. Hallett, P. M. Rossini, A. Pascual-Leone, and Safety of TMS Consensus Group, “Safety, ethical considerations, and application guidelines for the use of transcranial magnetic stimulation in clinical practice and research,” Clin Neurophysiol, vol. 120, no. 12, pp. 2008–2039, 2009.

[43] P. Kundu, N. D. Brenowitz, V. Voon, Y. Worbe, P. E. Vértes, S. J. Inati, Z. S. Saad, P. A. Bandettini, and E. T. Bullmore, “Integrated strategy for improving functional connectivity mapping using multiecho fMRI,” Proc Natl Acad Sci U S A, vol. 110, no. 40, pp. 16 187–16 192, 2013.

[44] P. Kundu, S. J. Inati, J. W. Evans, W. M. Luh, and P. A. Bandettini, “Differentiating BOLD and non-BOLD signals in fMRI time series using multi-echo EPI,” Neuroimage, vol. 60, no. 3, pp. 1759–1770, 2012.

[45] S. Whitfield-Gabrieli and A. Nieto-Castanon, “Conn: a functional connectivity toolbox for correlated and anticorrelated brain networks,” Brain Connect, vol. 2, no. 3, pp. 125–141, 2012.

[46] M. Brett, J. L. Anton, R. Valabregue, and J. B. Poline, “Region of interest analysis using an SPM toolbox,” vol. 16, no. 2, 497, 2002, available on CD-ROM in NeuroImage.

[47] G. K. Murray, P. R. Corlett, L. Clark, M. Pessiglione, A. D. Blackwell, G. Honey, P. B. Jones, E. T. Bullmore, T. W. Robbins, and P. C. Fletcher, “Substantia nigra/ventral tegmental reward prediction error disruption in psychosis,” Mol Psychiatry, vol. 13, no. 3, pp. 267–276, 2008.

[48] D. Martinez, M. Slifstein, A. Broft, O. Mawlawi, D. R. Hwang, Y. Huang, T. Cooper, L. Kegeles, E. Zarahn, A. Abi-Dargham, S. N. Haber, and M. Laruelle, “Imaging human mesolimbic dopamine transmission with positron emission tomography. part ii: amphetamine-induced dopamine release in the functional subdivisions of the striatum,” J Cereb Blood Flow Metab, vol. 23, no. 3, pp. 285–300, 2003.

[49] C. Rosazza, L. Minati, F. Ghielmetti, M. L. Mandelli, and M. G. Bruzzone, “Functional connectivity during resting-state functional MR imaging: study of the correspondence between independent component analysis and region-of-interest-based methods,” AJNR Am J Neuroradiol, vol. 33, no. 1, pp. 180–187, 2012.

[50] C. F. Beckmann and S. M. Smith, “Probabilistic independent component analysis for functional magnetic resonance imaging,” IEEE Trans Med Imaging, vol. 23, no. 2, pp. 137–152, 2004.

[51] V. Kiviniemi, J. H. Kantola, J. Jauhiainen, A. Hyvärinen, and O. Tervonen, “Independent component analysis of nondeterministic fMRI signal sources,” Neuroimage, vol. 19, no. 2 Pt 1, pp. 253–260, 2003.

[52] X. N. Zuo, C. Kelly, J. S. Adelstein, D. F. Klein, F. X. Castellanos, and M. P. Milham, “Reliable intrinsic connectivity networks: test-retest evaluation using ICA and dual regression approach,” Neuroimage, vol. 49, no. 3, pp. 2163–2177, 2010.

[53] T. E. Nichols and A. P. Holmes, “Nonparametric permutation tests for functional neuroimaging: a primer with examples,” Hum Brain Mapp, vol. 15, no. 1, pp. 1–25, 2002.

[54] I. Obeso, S. S. Cho, F. Antonelli, S. Houle, M. Jahanshahi, J. H. Ko, and A. P. Strafella, “Stimulation of the pre-SMA influences cerebral blood flow in frontal areas involved with inhibitory control of action,” Brain Stimul, vol. 6, no. 5, pp. 769–776, 2013.

[55] M. D. Fox, R. L. Buckner, M. P. White, M. D. Greicius, and A. Pascual-Leone, “Efficacy of transcranial magnetic stimulation targets for depression is related to intrinsic functional connectivity with the subgenual cingulate,” Biol Psychiatry, vol. 72, no. 7, pp. 595–603, 2012.

[56] A. Nordenskjöld, B. Mårtensson, A. Pettersson, E. Heintz, and M. Landén, “Effects of Hesel-coil deep transcranial magnetic stimulation for depression – a systematic review,” Nord J Psychiatry, vol. 70, no. 7, pp. 492–497, 2016.

[57] K. K. Kedzior, L. Gierke, H. M. Gellersen, and M. T. Berlim, “Cognitive functioning and deep transcranial magnetic stimulation (DTMS) in major psychiatric disorders: A systematic review,” J Psychiatr Res, vol. 75, pp. 107–115, 2016.

[58] M. Ceccanti, M. Inghilleri, M. L. Attilia, R. Raccah, M. Fiore, A. Zangen, and M. Ceccanti, “Deep TMS on alcoholics: effects on cortisolemia and dopamine pathway modulation. a pilot study,” Can J Physiol Pharmacol, vol. 93, no. 4, pp. 283–290, 2015.

[59] L. Dinur-Klein, P. Dannon, A. Hadar, O. Rosenberg, Y. Roth, M. Kotler, and A. Zangen, “Smoking cessation induced by deep repetitive transcranial magnetic stimulation of the prefrontal and insular cortices: a prospective, randomized controlled trial,” Biol Psychiatry, vol. 76, no. 9, pp. 742–749, 2014.

[60] F. Spagnolo, M. A. Volonté, M. Fichera, R. Chieffo, E. Houdayer, M. Bianco, E. Coppi, A. Nuara, L. Straffi, G. Di Maggio, L. Ferrari, D. Dalla Libera, S. Velikova, G. Comi, A. Zangen, and L. Leocani, “Excitatory deep repetitive transcranial magnetic stimulation with H-coil as add-on treatment of motor symptoms in Parkinson’s disease: an open label, pilot study,” Brain Stimul, vol. 7, no. 2, pp. 297–300, 2014.

[61] G. Kranz, E. A. Shamim, P. T. Lin, G. S. Kranz, and M. Hallett, “Transcranial magnetic brain stimulation modulates blepharospasm: a randomized controlled study,” Neurology, vol. 75, no. 16, pp. 1465–1471, 2010.

[62] C. Rapinesi, A. Del Casale, P. Scatena, G. D. Kotzalidis, S. Di Pietro, V. R. Ferri, F. S. Bersani, R. Brugnoli, R. N. Raccah, A. Zangen, S. Ferracuti, F. Orzi, P. Girardi, and G. Sette, “Add-on deep transcranial magnetic stimulation (dTMS) for the treatment of chronic migraine: A preliminary study,” Neurosci Lett, vol. 623, pp. 7–12, 2016.

[63] P. S. Tofts, “The distribution of induced currents in magnetic stimulation of the nervous system,” Phys Med Biol, vol. 35, no. 8, pp. 1119–1128, 1990.

[64] M. G. Stokes, C. D. Chambers, I. C. Gould, T. R. Henderson, N. E. Janko, N. B. Allen, and J. B. Mattingley, “Simple metric for scaling motor threshold based on scalp–cortex distance: application to studies using transcranial magnetic stimulation,” J Neurophysiol, vol. 94, no. 6, pp. 4520–4527, 2005.

[65] P. Trillenberg, S. Bremer, S. Oung, C. Erdmann, A. Schweikard, and L. Richter, “Variation of stimulation intensity in transcranial magnetic stimulation with depth,” J Neurosci Methods, vol. 211, no. 2, pp. 185–190, 2012.

[66] J. P. Brasil-Neto, L. G. Cohen, M. Panizza, J. Nilsson, B. J. Roth, and M. Hallett, “Optimal focal transcranial magnetic activation of the human motor cortex: effects of coil orientation, shape of the induced current pulse, and stimulus intensity,” J Clin Neurophysiol, vol. 9, no. 1, pp. 132–136, 1992.

[67] P. T. Fox, S. Narayana, N. Tandon, H. Sandoval, S. P. Fox, P. Kochunov, and J. L. Lancaster, “Column-based model of electric field excitation of cerebral cortex,” Hum Brain Mapp, vol. 22, no. 1, pp. 1–14, 2004.

[68] D. Balslev, W. Braet, C. McAllister, and R. C. Miall, “Inter-individual variability in optimal current direction for transcranial magnetic stimulation of the motor cortex,” J Neurosci Methods, vol. 162, no. 1–2, pp. 309–313, 2007.

[69] A. Opitz, M. Windhoff, R. M. Heidemann, R. Turner, and A. Thielscher, “How the brain tissue shapes the electric field induced by transcranial magnetic stimulation,” Neuroimage, vol. 58, no. 3, pp. 849–859, 2011.

[70] A. Thielscher, A. Opitz, and M. Windhoff, “Impact of the gyral geometry on the electric field induced by transcranial magnetic stimulation,” Neuroimage, vol. 54, no. 1, pp. 234–243, 2011.

[71] J. D. Bijsterbosch, A. T. Barker, K. H. Lee, and P. W. Woodruff, “Where does transcranial magnetic stimulation (TMS) stimulate? modelling of induced field maps for some common cortical and cerebellar targets,” Med Biol Eng Comput, vol. 50, no. 7, pp. 671–681, 2012.

[72] A. M. Janssen, T. F. Oostendorp, and D. F. Stegeman, “The coil orientation dependency of the electric field induced by TMS for M1 and other brain areas,” J Neuroeng Rehabil, vol. 12, no. 47, 2015.

[73] A. M. Janssen, S. M. Rampersad, F. Lucka, B. Lanfer, S. Lew, U. Aydin, C. H. Wolters, D. F. Stegeman, and T. F. Oostendorp, “The influence of sulcus width on simulated electric fields induced by transcranial magnetic stimulation,” Phys Med Biol, vol. 58, no. 14, pp. 4881–4896, 2013.

[74] A. M. Janssen, T. F. Oostendorp, and D. F. Stegeman, “The effect of local anatomy on the electric field induced by TMS: evaluation at 14 different target sites,” Med Biol Eng Comput, vol. 52, no. 10, pp. 873–883, 2014.

[75] M. Lu and S. Ueno, “Calculating the induced electromagnetic fields in real human head by deep transcranial magnetic stimulation,” in Conf Proc IEEE Eng Med Biol Soc, 2013, pp. 795–798.

[76] Z. D. Deng, S. H. Lisanby, and A. V. Peterchev, “Coil design considerations for deep transcranial magnetic stimulation,” Clin Neurophysiol, vol. 125, no. 6, pp. 1202–1212, 2014.

[77] Z. D. Deng, S. H. Lisanby, and A. V. Peterchev, “Electric field depth–focality tradeoff in transcranial magnetic stimulation: simulation comparison of 50 coil designs,” Brain Stimul, vol. 6, no. 1, pp. 1–13, 2013.

[78] T. Kammer, S. Beck, M. Erb, and W. Grodd, “The influence of current direction on phosphene thresholds evoked by transcranial magnetic stimulation,” Clin Neurophysiol, vol. 112, no. 11, pp. 2015–2021, 2001.

[79] S. Bestmann, J. Baudewig, H. R. Siebner, J. C. Rothwell, and J. Frahm, “BOLD MRI responses to repetitive TMS over human dorsal premotor cortex,” Neuroimage, vol. 28, no. 1, pp. 22–29, 2005.

[80] S. Bestmann, J. Baudewig, H. R. Siebner, J. C. Rothwell, and J. Frahm, “Functional MRI of the immediate impact of transcranial magnetic stimulation on cortical and subcortical motor circuits,” Eur J Neurosci, vol. 19, no. 7, pp. 1950–1962, 2004.

[81] S. Bestmann, C. C. Ruff, F. Blankenburg, N. Weiskopf, J. Driver, and J. C. Rothwell, “Mapping causal interregional influences with concurrent TMS–fMRI,” Exp Brain Res, vol. 191, no. 4, pp. 383–402, 2008.

[82] A. P. Strafella, J. H. Ko, J. Grant, M. Fraraccio, and O. Monchi, “Corticostriatal functional interactions in Parkinson’s disease: a rTMS/[11C]raclopride PET study,” Eur J Neurosci, vol. 22, no. 11, pp. 2946–2952, 2005.

[83] A. T. Sack, A. Kohler, D. E. Linden, R. Goebel, and L. Muckli, “The temporal characteristics of motion processing in hMT/V5+: combining fMRI and neuronavigated TMS,” Neuroimage, vol. 29, no. 4, pp. 1326–1335, 2006.

[84] H. R. Siebner, T. O. Bergmann, S. Bestmann, M. Massimini, H. Johansen-Berg, H. Mochizuki, D. E. Bohning, E. D. Boorman, S. Groppa, C. Miniussi, A. Pascual-Leone, R. Huber, P. C. Taylor, R. J. Ilmoniemi, L. De Gennaro, A. P. Strafella, S. Kähkönen, S. Klöppel, G. B. Frisoni, M. S. George, M. Hallett, S. A. Brandt, M. F. Rushworth, U. Ziemann, J. C. Rothwell, N. Ward, L. G. Cohen, J. Baudewig, T. Paus, Y. Ugawa, and P. M. Rossini, “Consensus paper: combining transcranial stimulation with neuroimaging,” Brain Stimul, vol. 2, no. 2, pp. 58–80, 2009.

[85] A. M. Speer, B. E. Benson, T. K. Kimbrell, E. M. Wassermann, M. W. Willis, P. Herscovitch, and R. M. Post, “Opposite effects of high and low frequency rTMS on mood in depressed patients: relationship to baseline cerebral activity on PET,” J Affect Disord, vol. 115, no. 3, pp. 386–394, 2009.

[86] I. Sibon, A. P. Strafella, P. Gravel, J. H. Ko, L. Booij, J. P. Soucy, M. Leyton, M. Diksic, and C. Benkelfat, “Acute prefrontal cortex TMS in healthy volunteers: effects on brain 11C-alphaMtrp trapping,” Neuroimage, vol. 34, no. 4, pp. 1658–1664, 2007.

[87] J. H. Ko, O. Monchi, A. Ptito, P. Bloomfield, S. Houle, and A. P. Strafella, “Theta burst stimulation-induced inhibition of dorsolateral prefrontal cortex reveals hemispheric asymmetry in striatal dopamine release during a set-shifting task: a TMS-[(11)C]raclopride PET study,” Eur J Neurosci, vol. 28, no. 10, pp. 2147–2155, 2008.

[88] A. M. Speer, T. A. Kimbrell, E. M. Wassermann, J. D. Repella, M. W. Willis, P. Herscovitch, and R. M. Post, “Opposite effects of high and low frequency rTMS on regional brain activity in depressed patients,” Biol Psychiatry, vol. 48, no. 12, pp. 1133–1141, 2000.

[89] R. Chen, J. Classen, C. Gerloff, P. Celnik, E. M. Wassermann, M. Hallett, and L. G. Cohen, “Depression of motor cortex excitability by low-frequency transcranial magnetic stimulation,” Neurology, vol. 48, no. 5, pp. 1398–1403, 1997.

[90] E. M. Wassermann, J. Grafman, C. Berry, C. Hollnagel, K. Wild, K. Clark, and M. Hallett, “Use and safety of a new repetitive transcranial magnetic stimulator.” Electroencephalogr Clin Neurophysiol, vol. 101, no. 5, pp. 412–417, 1996.

[91] W. Gerschlager, H. R. Siebner, and J. C. Rothwell, “Decreased corticospinal excitability after subthreshold 1 Hz rTMS over lateral premotor cortex,” Neurology, vol. 57, no. 3, pp. 449–455, 2001.

[92] L. H. Strens, A. Oliviero, B. R. Bloem, W. Gerschlager, J. C. Rothwell, and P. Brown, “The effects of subthreshold 1 Hz repetitive TMS on cortico-cortical and interhemispheric coherence,” Clin Neurophysiol, vol. 113, no. 8, pp. 1279–1285, 2002.

[93] K. Rubia, “Functional brain imaging across development,” Eur Child Adolesc Psychiatry, vol. 22, no. 12, pp. 719–731, 2013.

[94] J. S. Provost, A. Hanganu, and O. Monchi, “Neuroimaging studies of the striatum in cognition Part I: healthy individuals,” Front Syst Neurosci, vol. 9, no. 140, 2015.

[95] G. Bush, P. Luu, and M. I. Posner, “Cognitive and emotional influences in anterior cingulate cortex,” Trends Cogn Sci, vol. 4, no. 6, pp. 215–222, 2000.

[96] M. M. Botvinick, J. D. Cohen, and C. S. Carter, “Conflict monitoring and anterior cingulate cortex: an update,” Trends Cogn Sci, vol. 8, no. 12, pp. 539–546, 2004.

[97] C. S. Li and R. Sinha, “Inhibitory control and emotional stress regulation: neuroimaging evidence for frontal-limbic dysfunction in psycho-stimulant addiction,” Neurosci Biobehav Rev, vol. 32, no. 3, pp. 581–597, 2008.

[98] C. D. Chambers, H. Garavan, and M. A. Bellgrove, “Insights into the neural basis of response inhibition from cognitive and clinical neuroscience,” Neurosci Biobehav Rev, vol. 33, no. 5, pp. 631–646, 2009.

[99] W. Schultz, P. Dayan, and P. R. Montague, “A neural substrate of prediction and reward,” Science, vol. 275, no. 5306, pp. 1593–1599, 1997.

[100] L. H. Corbit, J. L. Muir, and B. W. Balleine, “The role of the nucleus accumbens in instrumental conditioning: Evidence of a functional dissociation between accumbens core and shell,” J Neurosci, vol. 21, no. 9, pp. 3251–3260, 2001.

[101] M. Pessiglione, B. Seymour, G. Flandin, R. J. Dolan, and C. D. Frith, “Dopamine-dependent prediction errors underpin reward-seeking behaviour in humans,” Nature, vol. 442, no. 7106, pp. 1042–1045, 2006.

[102] B. J. Harrison, C. Soriano-Mas, J. Pujol, H. Ortiz, M. López-Solà, R. HernándezRibas, J. Deus, P. Alonso, M. Yücel, C. Pantelis, J. M. Menchon, and N. Cardoner, “Altered corticostriatal functional connectivity in obsessive-compulsive disorder,” Arch Gen Psychiatry, vol. 66, no. 11, pp. 1189–1200, 2009.

[103] L. Cocchi, B. J. Harrison, J. Pujol, I. H. Harding, A. Fornito, C. Pantelis, and M. Yücel, “Functional alterations of large-scale brain networks related to cognitive control in obsessive-compulsive disorder,” Hum Brain Mapp, vol. 33, no. 5, pp. 1089–1106, 2012.

[104] A. Anticevic, S. Hu, S. Zhang, A. Savic, E. Billingslea, S. Wasylink, G. Repovs, M. W. Cole, S. Bednarski, J. H. Krystal, M. H. Bloch, C. S. Li, and C. Pittenger, “Global resting-state functional magnetic resonance imaging analysis identifies frontal cortex, striatal, and cerebellar dysconnectivity in obsessive-compulsive disorder,” Biol Psychiatry, vol. 75, no. 8, pp. 595–605, 2014.

[105] R. Z. Goldstein and N. D. Volkow, “Dysfunction of the prefrontal cortex in addiction: neuroimaging findings and clinical implications,” Nat Rev Neurosci, vol. 12, no. 11, pp. 652–669, 2011.

[106] S. Kühn and J. Gallinat, “Common biology of craving across legal and illegal drugs – a quantitative meta-analysis of cue-reactivity brain response,” Eur J Neurosci, vol. 33, no. 7, pp. 1318–1326, 2011.

[107] J. L. Abelson, G. C. Curtis, O. Sagher, R. C. Albucher, M. Harrigan, S. F. Taylor, B. Martis, and B. Giordani, “Deep brain stimulation for refractory obsessive-compulsive disorder,” Biol Psychiatry, vol. 57, no. 5, pp. 510–516, 2005.

[108] L. Mallet, M. Polosan, N. Jaafari, N. Baup, M. L. Welter, D. Fontaine, S. T. du Montcel, J. Yelnik, I. Chéreau, C. Arbus, S. Raoul, B. Aouizerate, P. Damier, S. Chabardès, V. Czernecki, C. Ardouin, M. O. Krebs, E. Bardinet, P. Chaynes, P. Burbaud, P. Cornu, P. Derost, T. Bougerol, B. Bataille, V. Mattei, D. Dormont, B. Devaux, M. Vérin, J. L. Houeto, P. Pollak, A. L. Benabid, Y. Agid, P. Krack, B. Millet, A. Pelissolo, and STOC Study Group, “Subthalamic nucleus stimulation in severe obsessive-compulsive disorder,” N Engl J Med, vol. 359, no. 20, pp. 2121–2134, 2008.

[109] M. Figee, J. Luigjes, R. Smolders, C. E. Valencia-Alfonso, G. van Wingen, B. de Kwaasteniet, M. Mantione, P. Ooms, P. de Koning, N. Vulink, N. Levar, L. Droge, P. van den Munckhof, P. R. Schuurman, A. Nederveen, W. van den Brink, A. Mazaheri, M. Vink, and D. Denys, “Deep brain stimulation restores frontostriatal network activity in obsessive-compulsive disorder,” Nat Neurosci, vol. 16, no. 4, pp. 386–387, 2013.

[110] J. P. O’Reardon, H. B. Solvason, P. G. Janicak, S. Sampson, K. E. Isenberg, Z. Nahas, W. M. McDonald, D. Avery, P. B. Fitzgerald, C. Loo, M. A. Demitrack, M. S. George, and H. A. Sackeim, “Efficacy and safety of transcranial magnetic stimulation in the acute treatment of major depression: a multisite randomized controlled trial,” Biol Psychiatry, vol. 62, no. 11, pp. 1208–1216, 2007.

[111] P. S. Sachdev, C. K. Loo, P. B. Mitchell, T. F. McFarquhar, and G. S. Malhi, “Repetitive transcranial magnetic stimulation for the treatment of obsessive compulsive disorder: a double-blind controlled investigation,” Psychol Med, vol. 37, no. 11, pp. 1645–1649, 2007.

[112] K. Dunlop, B. Woodside, M. Olmsted, P. Colton, P. Giacobbe, and J. Downar, “Reductions in cortico-striatal hyperconnectivity accompany successful treatment of obsessive-compulsive disorder with dorsomedial prefrontal rTMS,” Neuropsychopharmacology, vol. 41, no. 5, pp. 1395–1403, 2016.

[113] A. Mantovani, S. H. Lisanby, F. Pieraccini, M. Ulivelli, P. Castrogiovanni, and S. Rossi, “Repetitive transcranial magnetic stimulation (rTMS) in the treatment of obsessive-compulsive disorder (OCD) and Tourette’s syndrome (TS),” Int J Neuropsychopharmacol, vol. 9, no. 1, pp. 95–100, 2006.

[114] A. Mantovani, H. B. Simpson, B. A. Fallon, S. Rossi, and S. H. Lisanby, “Randomized sham-controlled trial of repetitive transcranial magnetic stimulation in treatment-resistant obsessive-compulsive disorder,” Int J Neuropsychopharmacol, vol. 13, no. 2, pp. 217–227, 2010.

[115] Y. Bloch, S. Arad, and Y. Levkovitz, “Deep TMS add-on treatment for intractable Tourette syndrome: A feasibility study,” World J Biol Psychiatry, vol. 17, no. 7, pp. 557–561, 2016.

[116] U. Ziemann, “TMS induced plasticity in human cortex,” Rev Neurosci, vol. 15, no. 4, pp. 253–266, 2004.

[117] V. Di Lazzaro, F. Pilato, M. Dileone, P. Profice, A. Oliviero, P. Mazzone, A. Insola, F. Ranieri, P. A. Tonali, and J. C. Rothwell, “Low-frequency repetitive transcranial magnetic stimulation suppresses specific excitatory circuits in the human motor cortex,” J Physiol, vol. 586, no. 18, pp. 4481–4487, 2008.

[118] Z. D. Deng, S. H. Lisanby, and A. V. Peterchev, “Electric field strength and focality in electroconvulsive therapy and magnetic seizure therapy: a finite element simulation study,” J Neural Eng, vol. 8, no. 1, 016007, 2011.

